# Blast fungus unconventional effector secretion sorting requires pre-Golgi compartment protein MoGrh1

**DOI:** 10.64898/2026.10.06.757164

**Authors:** Nawaraj Dulal, Richard A. Wilson

## Abstract

Brefeldin A (BFA)-resistant unconventional protein secretion (UPS) is documented in cellular systems across taxa and has roles in disease, but mechanistic details remain sparse. The blast fungus *Magnaporthe oryzae* uses Golgi-bypass UPS to deploy plant immunity-suppressing cytoplasmic effector proteins into host rice cells to establish infection. However, few *M. oryzae* UPS components are known, and connections to other cellular processes are undefined. Here, using live-cell imaging of *M. oryzae* mutant strains, we show that MoGrh1, a resident of the pre-Golgi, BFA-resistant ER-Golgi intermediate compartment (ERGIC), is required for UPS cargo sorting under the autophagy-inducing conditions of early host cell colonization. The loss of *MoGRH1* abolished UPS, with cytoplasmic effectors misrouted through a BFA-sensitive secretion pathway instead. Treatment with autophagy-suppressing α-ketoglutarate, a preferred carbon source, reconstituted unconventional secretion in Δ*grh1*, restoring fungal growth in host cells. We conclude that MoGrh1 supports UPS cargo sorting under the nutrient-limiting conditions of early host cell infection, linking the ERGIC and autophagy to disease-causing effector secretion in a devastating plant pathogen.

## Introduction

Proteins carrying leader N-terminal signal peptide (SP) sequences are often trafficked via the conventional ER-Golgi secretory route. Recognition particles bind SPs to direct translating cytosolic ribosomes to the ER, leading to peptide translocation, SP cleavage, and co-translational protein folding in the ER lumen. Folded proteins are packaged into coat protein II (COPII) vesicles for delivery to the early Golgi intermediate compartment (ERGIC). Further sorting and trafficking from the ERGIC to *cis*-Golgi, followed by intra-Golgi transport, is mediated by COPI vesicles, with cargos terminally released at the plasma membrane (PM) (Marie et al. 2008; Gomez-Navarro and Miller, 2016; Flood et al. 2025; Tojima and Nokana, 2026). COPI vesicles also mediate recycling from Golgi to ER. Conventional ER-Golgi secretion is inhibited by treatment with brefeldin A (BFA), which blocks the ARF GTPase activity regulating COPI vesicle biogenesis (Marie et al. 2008; Lin, 2025; Tojima and Nokano, 2026). BFA treatment collapses the Golgi, which is then absorbed into the ER, but the ERGIC retains its distinct structural identity and dynamics while undergoing positional changes (Marie et al 2008; Saraste and Marie, 2018). The ERGIC is a sorting compartment not only for classical secretion, but also for a subset of BFA-insensitive unconventional protein secretion (UPS) routes – the vesicular UPS pathways – which occur under both ER stress and physiological conditions, with trafficked proteins leaving the ERGIC and bypassing Golgi in intermediate vesicles (Lin, 2025; Saraste and Prydz, 2026; Zhang et al. 2026). Type III vesicular UPS involves delivery of cytosolic leaderless proteins directly to the ERGIC (Sun et al. 2024), whereas in Type IV UPS, cargo with SPs are delivered from ER lumen to the ERGIC in COPII vesicles (Ge et al. 2013; Rabouille, 2017; Saraste and Marie, 2018; Saraste and Prydz, 2026; Zhang et al. 2026). The intermediates involved in Golgi-independent, ERGIC-to-cell surface transport are largely undefined but include ERGIC-like compartments, autophagosomes and recycling endosomes (Saraste and Marie, 2018; Ge et al. 2013; Noh et al. 2018; Saraste and Prydz, 2026, Zhang et al. 2026). The ERGIC is also a membrane source for autophagosome biogenesis under cellular stress conditions, supplying ERGIC-derived COPII vesicles as phagophore membrane precursors (Ge et al. 2013; Ge et al. 2017; Gui et al. 2019; Saraste and Prydz, 2026). Depletion of the ERGIC or the specific ERGIC-resident proteins involved inhibits starvation-induced autophagosome formation (Ge et al. 2013; Ge et al. 2014; Ge et al. 2017; Li et al. 2021). Thus, the ERGIC has emerged as a central sorting hub – with ties to autophagy – involved in trafficking both classical and unconventional secretion cargo, but how this is precisely achieved is not fully defined (Saraste and Prydz, 2026).

In the devastating rice blast fungus *Magnaporthe oryzae* (Wilson and Talbot, 2009; Wilson, 2022; Valent, 2025), both conventional and unconventional protein secretion pathways are intrinsic to the host rice cell infection process (Giraldo et al. 2013; Dulal and Wilson, 2024). However, whether the ERGIC, which is a distinct compartment in yeast (Tojima et al. 2024), plays a role, and how sorting occurs, are unknown (Dulal and Wilson, 2024). Blast disease destroys up to one-third of rice harvests each year (Wilson and Talbot, 2009; Valent, 2025). Following host leaf penetration by specialized appressorial cells, *M. oryzae*, a filamentous fungus, elaborates bulbous, branching invasive hyphae (IH) that push into underlying epidermal cells, where IH become separated from living host cell cytoplasm by a host-derived-extra invasive hyphal membrane (EIHM) (Kankanala et al. 2007; Wilson and Talbot, 2009). IH fill this first infected living cell in a growth process driven by cycles of autophagy that liberate α-ketoglutarate to (re)activate TOR signaling (Li et al. 2023a) before moving at around 44 hours post inoculation (hpi) to adjacent, living cells. This biotrophic growth stage, which lasts for around 5 days before disease symptoms appear, is facilitated by the secretion of protein effectors – only expressed *in planta* – that mask the fungus from host detection (apoplastic effectors) or interfere with host innate immunity induction (cytoplasmic effectors) (Khang et al. 2010; Giraldo et al. 2013; Li et al. 2023b; Dulal and Wilson, 2024).

Apoplastic effectors are secreted into the matrix between the fungal cell wall and the EIHM by the conventional ER-Golgi pathway, with BFA treatment resulting in apoplastic effector internalization (Giraldo et al. 2013). In contrast, cytoplasmic effectors, which carry recognizable SPs, are first secreted unconventionally in a BFA-insensitive manner into the biotrophic interfacial complex (BIC), a specialized plant lipid-derived structure located outside IH, before translocation into host cells from the BIC via clathrin-mediated endocytosis (Giraldo et al. 2013; Oliveira-Garcia et al. 2023). Cytoplasmic effector secretion into the BIC occurs by the Type IV UPS vesicular pathway (Dulal and Wilson, 2024) and requires the exocyst complex and the t-SNARE Sso1 (Giraldo et al. 2013), which likely fuse secretory vesicles to the PM. Upstream of vesicle-PM fusion, at the ribosome-ER boundary, a tRNA modification and codon usage bias determines cytoplasmic effector mRNA translation efficiency (Li et al. 2023b). What lies between these extremes in the *M. oryzae* UPS pathway is unknown.

To better understand the cellular and molecular processes governing effector cargo sorting and trafficking in *M. oryzae*, we targeted the uncharacterized gene *MoGRH1*, an orthologue of the nonessential yeast gene *GRH1* (Behnia et al. 2007), encoding the sole relatives in their respective fungi of the mammalian Golgi Reassembly Stacking Protein of 55 kDa (GRASP55) and GRASP65 (Saraste and Prydz, 2026). Although GRASPs were first associated with Golgi reassembly and stacking, they also have roles in Type III and Type IV UPS vesicular pathways (Nuchel et al. 2021). In yeast (which, like other fungi, lack cisternal Golgi stacks), Grh1 was recently classified as an ERGIC (rather than *cis-*Golgi) marker protein (Tojima et al. 2024; Saraste and Prydz, 2026; Tojima and Nokana, 2026). Yeast Grh1 interacts with COPII vesicles (Behnia et al. 2007) and is involved in secretory autophagosome biogenesis but not degradative autophagy (Bruns et al. 2011). Grh1 also acts as a scaffold for vesicular interactions during UPS involving autophagosomes, endosomes and ERGIC compartments (Bruns et al. 2011; Tojima et al. 2024; Zhang et al. 2026), ultimately connecting the ERGIC with *trans*-Golgi recycling endosomes to bypass Golgi (Saraste and Prydz, 2026). Its loss in yeast, while tolerated, leads to less organized secretory compartments and impacts UPS but not ER-Golgi secretory pathways (Levi et al. 2010). Consistent with ERGIC residency, BFA treatment causes yeast Grh1 and other pre-/early-Golgi proteins to localize to puncta typical of BFA generated ERGIC compartments, in contrast to Golgi proteins that are absorbed into the ER as the Golgi is dismantled (Tojima and Nakano, 2026).

Here, we show that in *M. oryzae*, MoGrh1 resides in BFA-resistant ERGIC-like structures that associate with cytoplasmic effectors. The loss of MoGrh1 impaired biotrophic growth, abolished autophagic flux, and attenuated the UPS pathway (but not ER-Golgi secretion), resulting in the disorganized secretion of cytoplasmic effectors via the conventional, BFA-sensitive ER-Golgi secretion system. Treating *M. oryzae* infected rice cells with α-ketoglutarate (a preferred carbon source that suppresses autophagy in *M. oryzae* (Li et al. 2023a)) remediated biotrophic growth and UPS in Δ*grh1*. Autophagy induction with rapamycin also restored UPS in Δ*grh1*, although end-stage autophagic flux remained impaired. We conclude that effector deployment by UPS is constitutive, with ERGIC cargo sorting and pathway integrity critically supported by MoGrh1-dependent autophagic flux control under the nutrient-limiting conditions of early host cell infection. Together, our findings shed light on the rules of effector secretion in a devastating plant pathogen that may have implications for our understanding of unconventional secretion in other systems and contexts.

## Results

### *MoGRH1* is required for host rice cell colonization

Using the *MoGRH1* sequence at locus MGG_05757 in the *M. oryzae* genome (Dean et al. 2003), we used homologous recombination to replace 1 kb of the 5’ *MoGRH1* coding region in the wild type (WT) isolate Guy11 with the *ILV2* gene conferring sulphonyl urea resistance. After screening for resistance and confirming the gene deletion by PCR, we tested five independent transformants to ensure they behaved similarly with regards to plate growth, sporulation rates and infection capacity (examples shown in **Fig. 1A** and **Supplementary Fig. S1A,B**). We picked one of these Δ*grh1*-carrying mutant strains to receive the *GRH1* gene (fused to *eGFP*) and generate the Δ*grh1 GRH1-GFP* complementation strain. Δ*grh1* mutant strain colonies had marginally reduced colony diameters compared to WT and the Δ*grh1 GRH1-GFP* complementation strain after 10 days growth on plates of complete media (CM), but sporulation rates were significantly (one-way ANOVA followed by Tukey’s HSD, *P* < 0.05) reduced (**Supplementary Fig. S1A,B**). When equal numbers of spores were applied to three-week-old seedlings of the susceptible rice cultivar CO-39, Δ*grh1* strains were found to be nonpathogenic, producing only small flecks on leaves that failed to produce spores rather than the spreading necrotic lesions typical of WT and recapitulated in the complementation strain (**Fig. 1A**). Loss of pathogenicity by the Δ*grh1* mutant strain was not due to impaired appressorium formation or the inability of appressoria to penetrate host cells, whose rates were not significantly different (one-way ANOVA followed by Tukey’s HSD, *P* < 0.05) from WT (**Supplementary Fig. S1C,D**). However, live-cell imaging of optically clear rice leaf sheaths showed that Δ*grh1* IH growth in the first infected rice epidermal cell was attenuated at 44 hpi compared to WT (**Fig. 1B**), with Δ*grh1* IH only rarely spreading into adjacent cells (**Supplementary Fig. S1E**). We conclude that *MoGRH1* is an essential virulence determinant that, in addition to maximizing sporulation, is required, post-penetration, for establishing biotrophic growth in living host rice cells and thus for facilitating disease progression.

**Figure 1.**
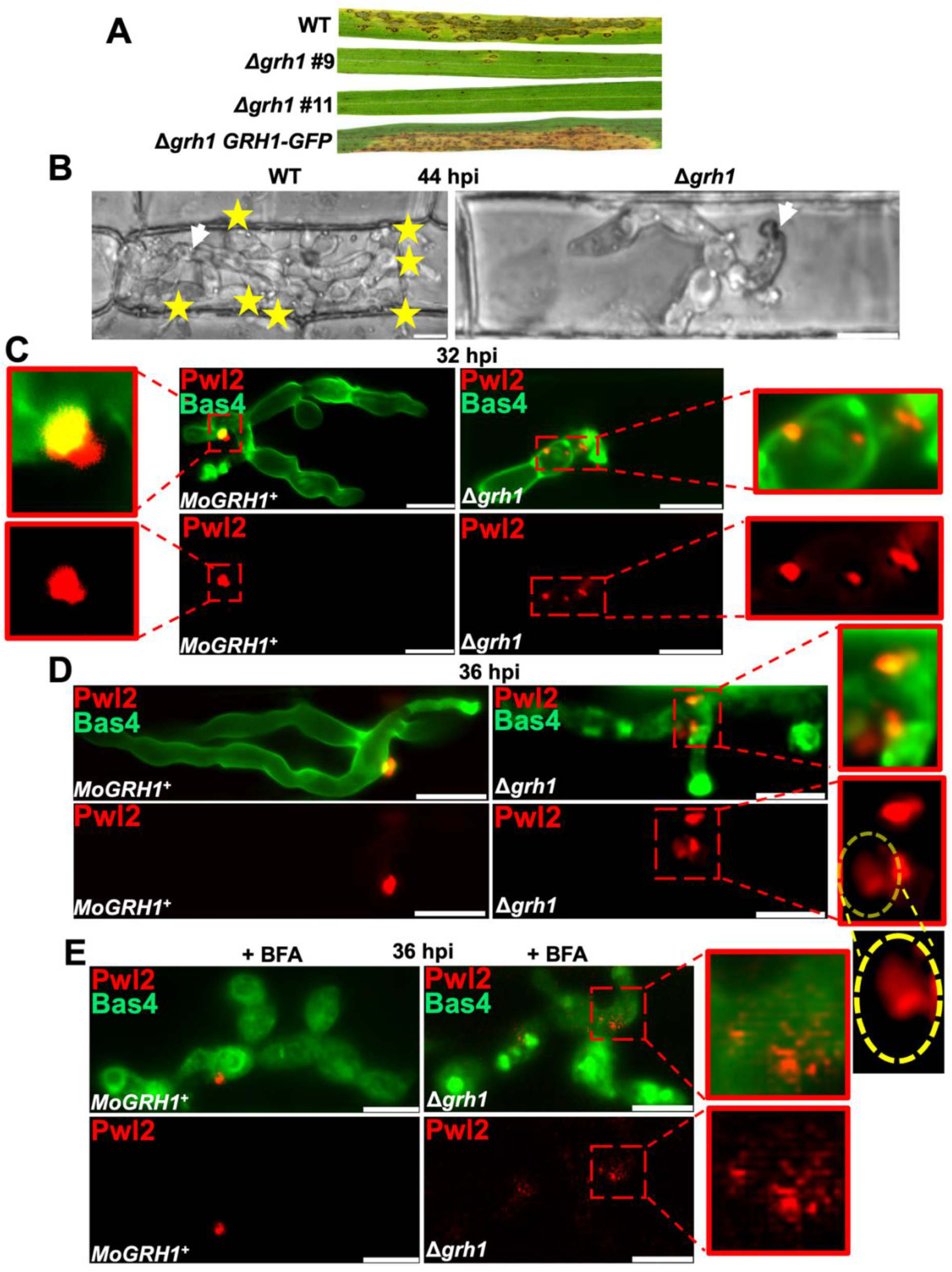
*MoGRH1* is required for biotrophic growth and unconventional effector secretion in host rice cells. **A.** The *M. oryzae MoGRH1* gene is required for full pathogenicity on leaves of the susceptible rice cultivar CO-39. Three-week-old CO-39 seedlings were spray-inoculated with spores of the indicated strains at a concentration of 1×10^5^ spores ml^-1^. Disease symptoms were photographed at 5 days post-inoculation. The images shown are representative of at least five leaves assessed in each of three independent biological replicates. **B.** Live-cell imaging of infected detached rice leaf sheath epidermal cells at 44 hpi showing that the Δ*grh1* strain is impaired in biotrophic growth compared to WT. White arrowheads mark appresorial penetration sites, yellow stars indicate IH that have spread into neighboring host cells. **C, D.** Live-cell imaging of detached rice leaf sheath epidermal cells at 32 hpi **(C)** and 36 hpi **(D)** following infection by the indicated strains producing BIC-localized Pwl2-mCherry:NLS and apoplast-localized Bas4-GFP. Hatched yellow circle highlights internalized, diffuse Pwl2 patches. **E.** Rice leaf-sheath cells were treated at 32 hpi with 50 µg ml^-1^ BFA in 0.1 % DMSO for 4 hr to inhibit conventional ER-Golgi secretion. In the *MoGRH1^+^*strain, BFA caused Bas4 retention in invasive hyphae but did not affect Pwl2 secretion into BICs, as expected. In the Δ*grh1* strain, BFA caused internalization of both Bas4 and Pwl2. **C-E.** Enlarged views of the indicated areas are shown in red boxes. Representative images were obtained by examining 50 infected rice cells per leaf sheath for each strain in three independent biological replicates. Scale bars are 10 μm.

### *MoGRH1* is required for effector sorting into the UPS pathway

To shed light on why *MoGRH1* was necessary for the *M. oryzae* biotrophic growth stage, we knocked out *MoGRH1* in our modified Guy11 strain that produces the BIC-localized cytoplasmic effector Pwl2 fused to mCherry:NLS, and the apoplastic effector Bas4 fused to eGFP (**Fig. 1C**). In the parental *MoGRH1^+^* strain, as previously reported (Khang et al. 2010; Giraldo et al. 2013; Marroquin-Guzman et al. 2017; Li et al. 2023a; Li et al. 2023b; Oliveira-Garcia et al. 2023), at 32 hpi and 36 hpi, Pwl2 localized to single punctate BICs in each primary infected rice cell, with Pwl2 accumulating in both BIC base- and outer layers, while Bas4 outlined IH in the apoplastic space and accumulated in the BIC base layer only (**Fig. 1C**). In contrast, in the Δ*grh1* background, Pwl2 accumulated in fragmented BICs external to IH at 32 hpi and, by 36 hpi, also to smaller diffuse intracellular patches (**Fig. 1C**). Therefore, the loss of *MoGRH1* negatively impacted focused Pwl2 deployment into the BIC.

Considering the documented role of yeast Grh1 in unconventional secretion, we hypothesized that the Δ*grh1* mutant strain misregulated Pwl2 deployment due to impaired UPS. To test this, we treated infected rice leaf sheaths with BFA – which inhibits COPI biosynthesis and collapses the Golgi into the ER (but does not destroy the ERGIC) – at 32 hpi and observed the effects by 36 hpi. In the *MoGRH1^+^*strain, as previously reported (Giraldo et al. 2013; Li et al. 2023b; Dulal and Wilson, 2024), Pwl2 secretion into the BIC was unaffected by BFA treatment, but Bas4 became internalized, indicating disruption to the conventional ER-Golgi secretion pathway (**Fig. 1D**). In the Δ*grh1* strain, however, in addition to Bas4, Pwl2 was also internalized following BFA treatment. The internalization of Pwl2 throughout IH following BFA treatment in Δ*grh1* was clearly seen in 3D side-view projections (**Fig. 2A-C**). We conclude that *MoGRH1* is required for cargo sorting and maintaining trafficking through the UPS pathway, but in the absence of *MoGRH1*, cargo is misrouted via a BFA-sensitive secretory process instead, leading to disorganized secretion into multiple external BIC-like structures. This pathway switch has not been described before for any previous *M. oryzae* mutant strain or other phytopathogen, nor have factors involved in UPS cargo sorting been previously deduced in *M. oryzae* (Dulal and Wilson, 2024).

**Figure 2.**
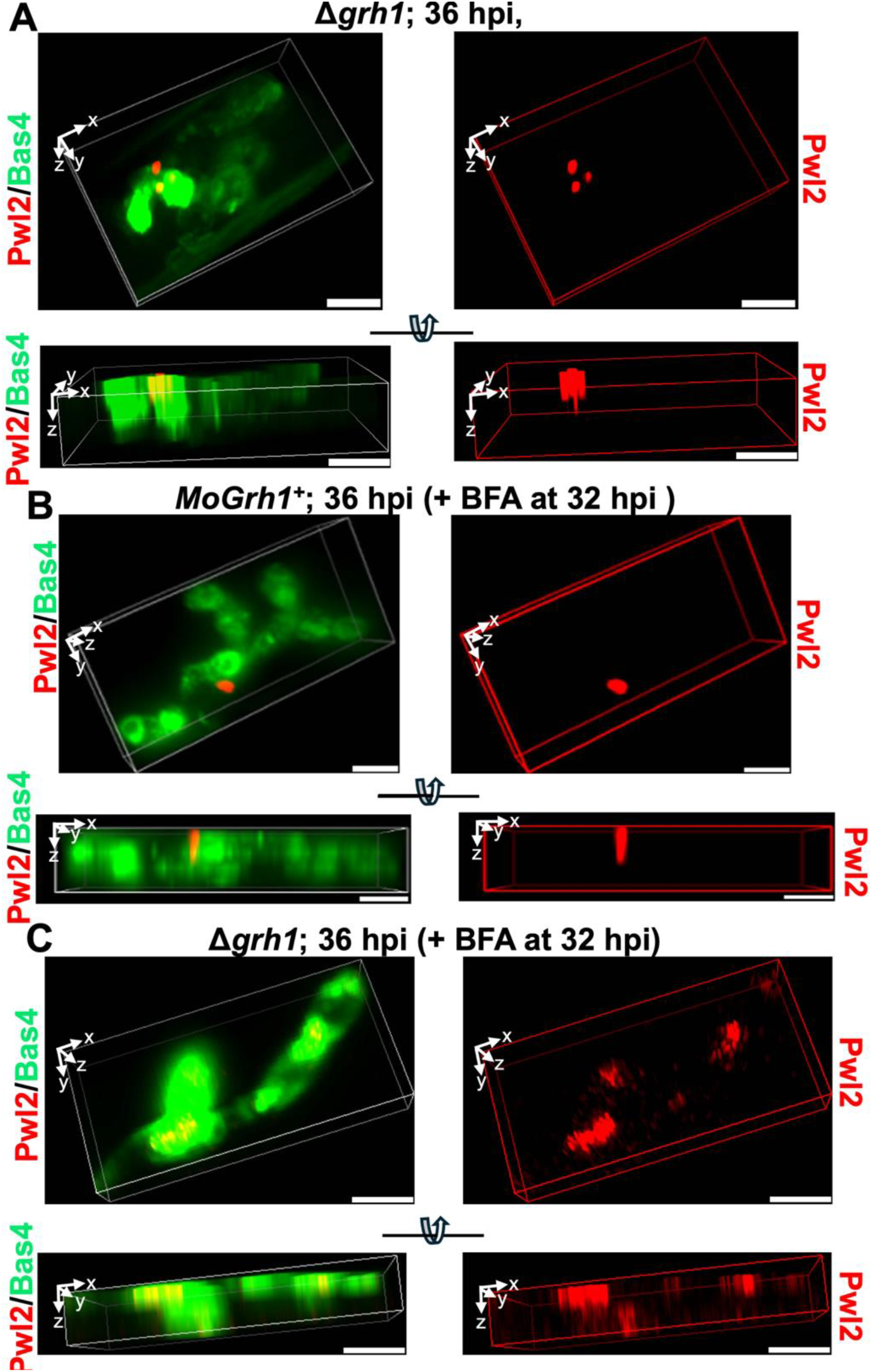
Loss of *MoGRH1* causes cytoplasmic effector secretion in a BFA-sensitive manner. **A-C.** Three-dimensional rendering from live-cell images of the indicated strains, at 36 hpi, expressing the BIC-localized effector Pwl2-mCherry:NLS and the apoplast-localized effector Bas4-GFP. For each treatment, the upper panels show a three-dimensional reconstruction tilted along the x-axis to facilitate visualization of BIC positions. The lower panels present the same reconstruction rotated to provide a basal, lateral view along the z-axis. x, y, and z indicate the axes of the micrographs. Representative images were selected from 50 infected rice cells examined per leaf sheath for each strain across three independent biological replicates. Scale bars =10 μm. **B,C.** Strains were treated at 32 hpi with 50 μg ml^-1^ BFA in 0.1 % DMSO and imaged at 36 hpi.

### MoGrh1 localizes to BFA-resistant compartments in invasive hyphae in rice cells

To better understand how *MoGRH1* maintained cytoplasmic effector secretion, we next studied the subcellular localization of MoGrh1 in IH *in planta*, finding that MoGrh1-GFP localization was dynamic as the biotrophic growth phase progressed. MoGrh1-GFP resided in a network of tubular structures that contacted vacuoles at 28 – 36 hpi, formed distinct puncta around vacuoles at 40 hpi, then became more uniform in the cytoplasm at 42 – 44 hpi as IH moved to adjacent cells (examples in **Fig. 3A** (36 hpi, 40 hpi and 42 hpi) and **Supplementary Fig. S2** (28 hpi, 32 hpi and 44 hpi)). Moreover, like the yeast ERGIC protein Grh1 but unlike proteins of the mature Golgi (Tojima et al. 2024; Tojima and Nakano, 2026), MoGrh1 was not absorbed into the ER following BFA treatment of IH in rice cells, instead persisting in punctate structures that contacted vacuoles and resembled the BFA-induced ERGIC aggregates in yeast, plant and vertebrate cells (Tojima et al. 2024; Saraste and Prydz, 2026; Tojima and Nakano, 2026) (**Fig. 3B**). We conclude that, consistent with the behavior of Grh1 in yeast, MoGrh1 in IH localizes to distinct, dynamic ERGIC-like structures that are stable under BFA treatment. This localization is in line with both the requirement of MoGrh1 for BFA-insensitive UPS in *M. oryzae* shown here, and with the ERGIC being involved in sorting unconventional as well as conventionally secreted cargo, as described in other systems (Saraste and Prydz, 2026).

**Figure 3.**
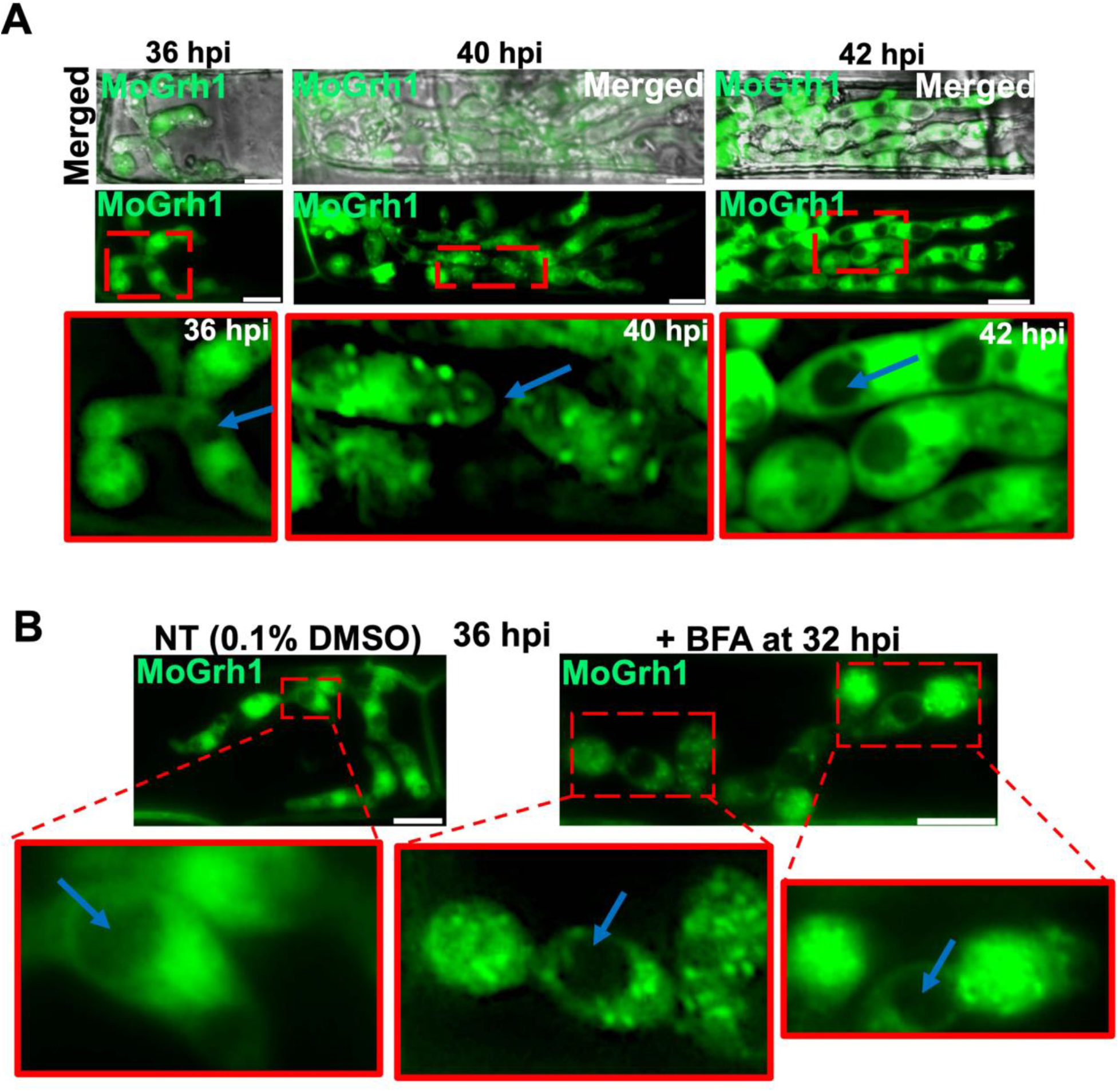
MoGrh1 localizes to BFA-resistant compartments. **A.** Live-cell imaging of the *Δgrh1 MoGRH1-GFP* complementation strain in detached rice leaf-sheath epidermal cells at the indicated time points showing dynamic changes in MoGrh1-GFP localization as biotrophy progresses, with MoGrh1 shifting from tubular cytoplasmic structures to punctate compartments before becoming more uniformly distributed around vacuoles. Images in solid red boxes are enlarged views of the area indicated in the dashed red boxes immediately above. **B.** Live-cell imaging of the Δ*grh1 MoGRH1-GFP* complementation strain at 36 hpi on detached rice leaf sheaths following treatment at 32 hpi with either 50 μg ml^-1^ BFA in 0.1 % DMSO or 0.1 % DMSO alone as the non-treated (NT) control. BFA treatment resulted in the retention of MoGrh1-GFP in ERGIC-like punctate structures. **A,B.** Blue arrows indicate examples of vacuoles. Representative images were selected from 50 infected rice cells examined per leaf sheath for each treatment, with each experiment repeated three times. Scale bars = 10 μm.

### Cytoplasmic effector Pwl2 traffics via BFA-induced ERGIC-like structures

The previous results revealed the existence of MoGrh1-associated, BFA-resistant ERGIC-like structures in *M. oryzae*. We hypothesized that *M. oryzae* cytoplasmic effectors were trafficked via this ERGIC sorting hub, and sought to test this by trapping a modified version of Pwl2 in BFA-induced ERGIC-like aggregates. We generated a Δ*grh1* complementation strain producing both MoGrh1-GFP and Pwl2-mCherry:HDEL. HDEL is the ER retention signal in yeast (KDEL in mammals) that interacts with H/KDEL-receptors in the ERGIC and Golgi to return escaped ER-resident proteins in COPI-coated vesicles (Fullekrug et al. 1997; Cabrera et al. 2003; Bruezza et al. 2004; Newstead and Barr, 2020; Tojima and Nakano, 2026). First, we confirmed that HDEL could route secreted Pwl2 back to the ER by analyzing Pwl2-HDEL and MoGrh1-GFP localization in IH in the absence of BFA. Under these conditions, Pwl2-mCherry:HDEL (which, like native Pwl2, carries a signal peptide sequence and is presumably trafficked out of the ER after co-translational folding) did not accumulate in the BIC like native Pwl2 and instead localized to tubular internal compartments distributed throughout IH, suggesting it was being actively returned to the ER by HDEL receptors (**Fig. 4A**). Pwl2-mCherry:HDEL-labelled compartments partially overlapped with MoGrh1-GFP, which is consistent with Grh1 in yeast, where a subset of Grh1-positive ERGIC puncta associate with ER exit sites (Bruns et al. 2011; Tojima et al. 2024; Tojima and Nakano, 2026). In contrast, instead of tubular structures, BFA treatment resulted in Pwl2-mCherry:HDEL accumulating mostly in large aggregates that overlapped more extensively with MoGrh1-GFP compared to the untreated control (**Fig. 4B**). When considering that BFA treatment in mammalian cells confines KDEL receptors and their KDEL-tagged protein ligands to the ERGIC (Fullekrug et al. 1997; Bruezza et al. 2004), our results taken together are consistent with Pwl2-mCherry:HDEL being trafficked back to the ER from the ERGIC in COPI vesicles such that BFA treatment (which disassembles the Golgi but not ER or the ERGIC) traps Pwl2-mCherry:HDEL in BFA-induced ERGIC aggregates. We conclude that to bypass Golgi, Pwl2 traffics via the ERGIC for MoGrh1-dependent UPS pathway sorting.

**Figure 4.**
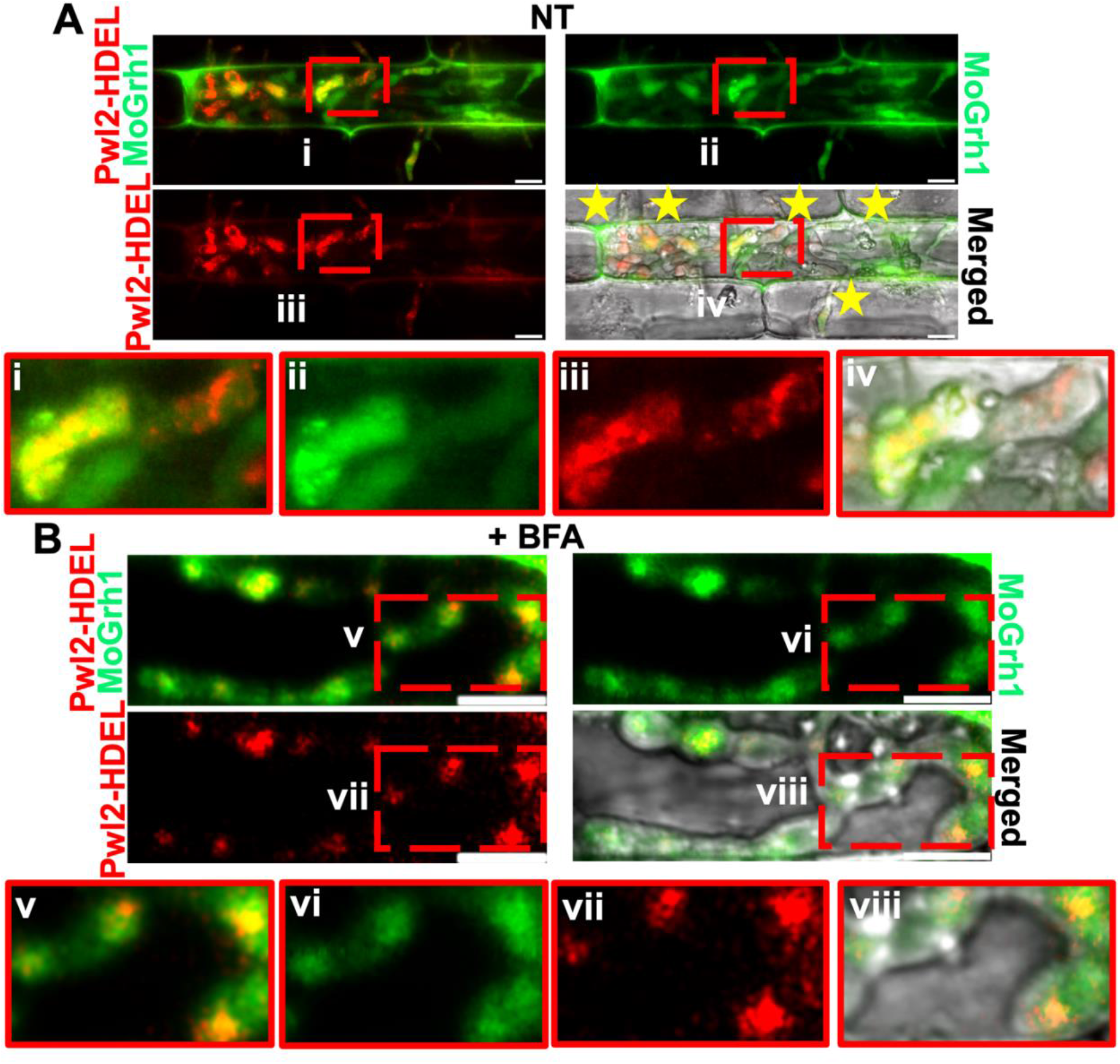
The unconventionally secreted cytoplasmic effector Pwl2 traffics via BFA-induced ERGIC compartments. **A.** Live-cell imaging at 42 hpi of detached rice leaf sheath cells inoculated with the indicated strain. Pwl2-HDEL refers to the Pwl2 protein carrying the C-terminal ER retention signal HDEL. Pwl2-mCherry-HDEL accumulated in vesicle-like compartments within IH and partially overlapped MoGrh1-GFP (yellow). Yellow asterisks mark IH that have extended into adjacent host cells. Enlarged views of the dashed, red-boxed regions marked i-iv are provided in solid red outlined boxes in the corresponding lower panels (i–iv). **B** Live-cell imaging performed at 36 hpi on detached rice leaf sheaths inoculated with the indicated strain following treatment at 32 hpi with 50 μg ml^-1^ BFA in 0.1 % DMSO. Enlarged views of the dashed, red-boxed regions marked v-viii are provided in solid red outlined boxes in the corresponding lower panels (v–viii). **A,B.** Representative images were selected from 50 infected rice cells examined per leaf sheath for each treatment, with each experiment repeated three times. Scale bars = 10 μm.

### *MoGRH1* is required for autophagic flux

Our next insight into the connection between *MoGRH1* and UPS came from determining *MoGRH1* was required for autophagic flux. A role for *MoGRH1* in autophagy was first suggested from plate tests showing how on nutrient-less water agar plates (our standard autophagy assay (Li et al. 2023a)), plugs of Δ*grh1* mycelia from CM plates transferred to water agar produced smaller colonies than WT or the complementation strain, indicating nutrient recycling deficiencies in the mutant (**Supplementary Fig. S3A**). Following up on this line of inquiry, we transformed WT and Δ*grh1* strains with a plasmid expressing *GFP-ATG8*. GFP-Atg8 is an autophagic flux marker (Klionsky et al. 2021), previously used by us to assess autophagy dynamics in *M. oryzae* vegetative hyphae and IH (Li et al. 2023a). Under autophagy-inactivating conditions (i.e. nutrient-rich CM media), GFP-Atg8 is cytoplasmic. Upon autophagy induction conditions (i.e. by switching from CM to H_2_O), GFP-Atg8 incorporates into autophagosomes which fuse to fungal vacuoles (lysosomes in animal cells), where GFP is cleaved and slowly degraded. Stable free green fluorescence in vacuoles/lysosomes signals end-stage autophagy and therefore is a measure of autophagic flux. Consistent with our previous reports (Li et al. 2023a), GFP-Atg8 in *MoGRH1^+^* mycelia was solely cytoplasmic following growth in nutrient-rich liquid CM, and the GFP signal shifted to vacuoles upon transfer into autophagy-inducing H_2_O (**Fig. 5A**). In Δ*grh1*, GFP-Atg8 was similarly observed in mycelial cytoplasm in CM media and was not localized to vacuoles. However, Δ*grh1* GFP-Atg8 often formed cytoplasmic puncta in CM. In H_2_O, these cytoplasmic GFP-Atg8 puncta in Δ*grh1* became more prominent, but vacuoles remained unlabeled with GFP (**Fig. 5A**). Previously, we showed that α-ketoglutarate, derived from glutaminolysis, was an *M. oryzae* preferred carbon source and nutrient-sufficiency signal that suppressed autophagy– even in the absence of a nitrogen source – when applied exogenously as the cell permeable analog dimethyl α-ketoglutarate (DMKG) (Li et al. 2023a). Here, in H_2_O treated with DMKG, autophagy was suppressed in both *MoGRH1^+^* and Δ*grh1* strains, and the GFP-Atg8-labelled puncta in Δ*grh1* IH were not observed (**Fig. 5B**). In summary, under axenic growth conditions, Δ*grh1* is impaired in autophagic flux, forming abnormal GFP-labelled cytoplasmic compartments – likely representing failed autophagosome trafficking or fusion to the vacuole – that are dispersed when autophagy is suppressed by α-ketoglutarate (**Fig. 5C**). This role of MoGrh1 in end-stage autophagic flux is consistent with that of mammalian GRASP55, which acts as a membrane tether to enhance autophagosome-lysosome fusion and whose loss similarly leads to decreased autophagic flux (but increased autophagosome formation) under carbon starvation conditions compared to WT (Zhang et al. 2018).

**Figure 5.**
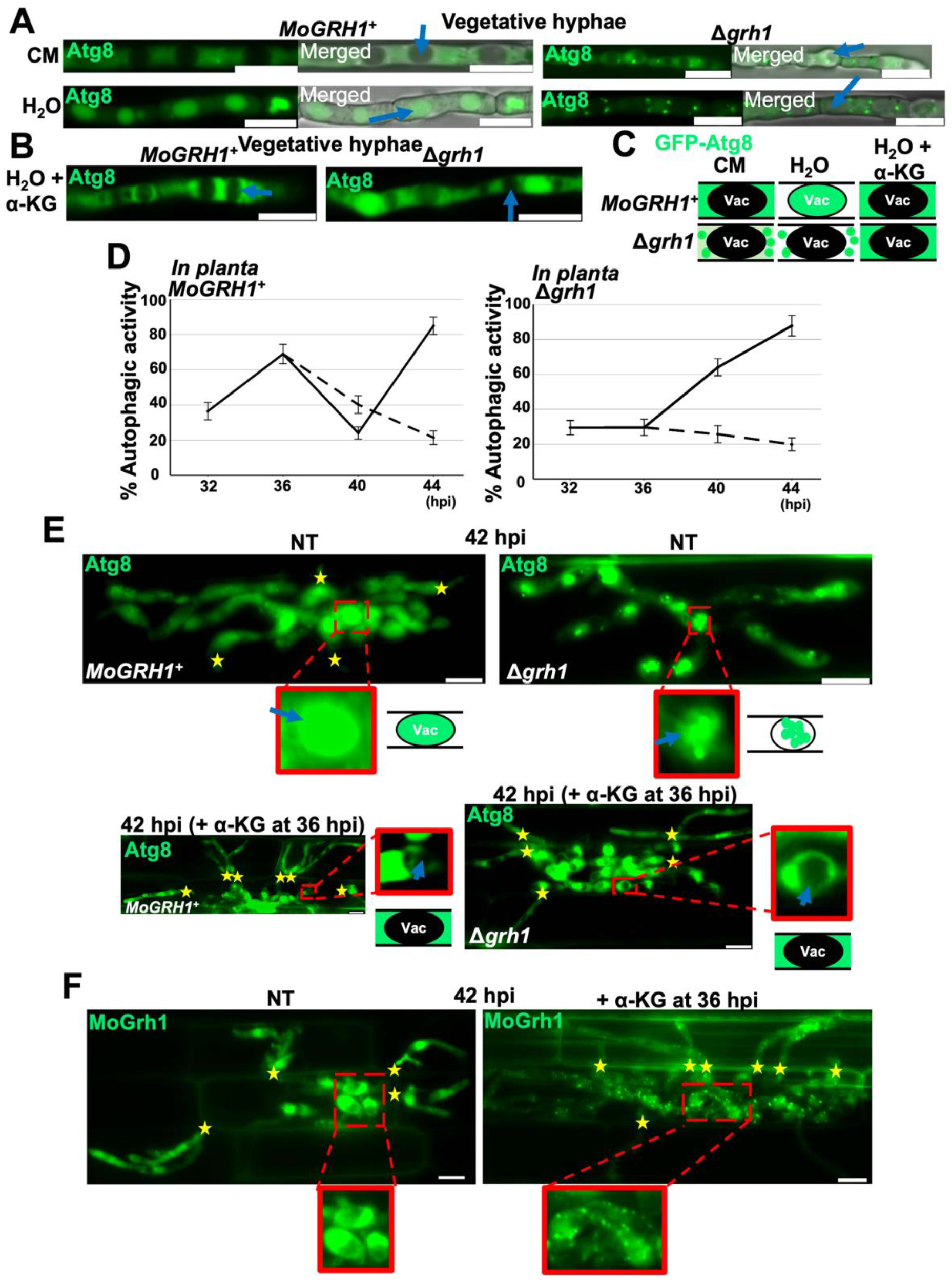
*MoGRH1* is required for autophagy. **A,B.** Micrographs of vegetative hyphae of the indicated strains. *GFP-ATG8*-expressing strains were cultured in liquid CM for 42 hr. Mycelia were then harvested, washed with water, and transferred to fresh liquid CM, water, or water containing 10 mM α-ketoglutarate (as DMKG), as indicated, for 4 hr before imaging. Blue arrows indicate examples of vacuoles. Images were selected from observing 50 mycelial fragments, repeated in triplicate. **C** Schematic summarizing autophagic activity in vegetative hyphae in (**A,B**). Vac denotes vacuoles. **D** Autophagic flux in the IH of the indicated strains was assessed *in planta* at the indicated time points. Solid lines represent untreated samples, dashed lines show autophagic activity at 40 and 44 hpi following treatment with 10 mM α-ketoglutarate (DMKG) at 36 hpi. Autophagic activity was calculated as the percentage of IH vacuoles exhibiting GFP accumulation in 50 infected rice cells examined per leaf sheath for each strain, in triplicate. Data represent the mean ± SD of three biological replicates. **E,F.** Live-cell imaging of detached rice leaf-sheath epidermal cells at 42 hpi following infection with the indicated strains expressing *GFP-ATG8* (**E**), or Δ*grh1* expressing *MoGRH1-GFP* (**F**). Samples were either untreated (NT) or treated with 10 mM α-ketoglutarate (DMKG) at 36 hpi. Yellow stars indicate invasive hyphae (IH) extending into neighboring host cells. Example vacuoles are marked with blue arrows. Representative images were selected from 50 infected rice cells examined per leaf sheath across three biological replicates. Scale bars = 10 μm.

Autophagic flux perturbation was observed in Δ*grh1* IH *in planta* compared to *MoGRH1^+^*strains (**Fig. 5D**; micrograph examples at each timepoint for both strains in **Supplementary Fig. S3B**). As described by us previously (Li et al. 2023a), autophagic flux in the *MoGRH1^+^*Guy11 strain producing GFP-Atg8 was cyclical during biotrophy, as measured by the percentage of vacuoles fluorescing green, in order to generate α-ketoglutarate to promote growth (**Fig. 5D**). In contrast, autophagic flux cycling was abolished in Δ*grh1* (quantified in **Fig. 5D**), with the GFP signal accumulating in vacuoles either weakly (at 40 hpi, **Supplementary Fig. S3B** - note that vacuoles were scored in **Fig. 5D** as completing autophagy if they fluoresced green regardless of fluorescence intensity), or aberrantly as puncta (at 42 hpi, **Fig. 5E**, and 44 hpi, **Supplementary Fig. S3B**). Such late-stage perturbed autophagic flux may be non-specific and/ or the result of alternative cellular stress pathways (Li et al. 2023a). Treating infected rice cells with DMKG at 36 hpi inhibited autophagy in both strains (**Fig. 5D**) and in Δ*grh1*, by 42 hpi, DMKG stimulated cell-to-cell movement (**Fig. 5E**). Furthermore, in the complementation strain, MoGrh1-GFP was distributed uniformly in the cytoplasm around vacuoles at 42 hpi when autophagy was active, but autophagy inactivation by DMKG treatment at 36 hpi resulted, by 42 hpi, in MoGrh1 localizing to puncta (**Fig. 5F**). Thus, MoGrh1 responds to the autophagy activation status of the cell and *MoGRH1* is required for autophagic cycling in IH to promote biotrophic growth. However, like other *M. oryzae* autophagy-related genes (Li et al. 2023a), *MoGRH1* is dispensable when autophagy is unneeded in the presence of exogenous α-ketoglutarate.

### α-ketoglutarate or rapamycin treatment restored UPS in Δ*grh1* IH in rice cells

We hypothesized that in addition to biotrophic growth, α-ketoglutarate treatment also remediated UPS and effector deployment in Δ*grh1* strains. We treated infected rice leaf sheaths with DMKG at 32 hpi and viewed them at 36 hpi (**Fig. 6A**) or treated them at 36 hpi and viewed them at 42 hpi (**Fig. 6B**). Compared to the untreated controls, at 36 hpi, a single punctate BIC in each infected rice cell was reconstituted by DMKG treatment in Δ*grh1* strains (**Fig. 6A**). At 42 hpi, DMKG treatment stimulated Δ*grh1* biotrophic growth into neighboring cells, and punctate BICs formed at the tips of each new IH (**Fig. 6B**), as previously observed for WT under native conditions (Khang et al. 2010; Sun et al. 2018; Li et al. 2023a). Remediated UPS pathway trafficking to the BIC in Δ*grh1* by DMKG was confirmed by the co-addition of BFA, which internalized Bas4 but not Pwl2 in DMKG + BFA treated Δ*grh1* IH (**Fig. 6C**). We conclude that exogenous α-ketoglutarate remediates UPS and restores focal BIC formation in Δ*grh1*.

**Figure 6.**
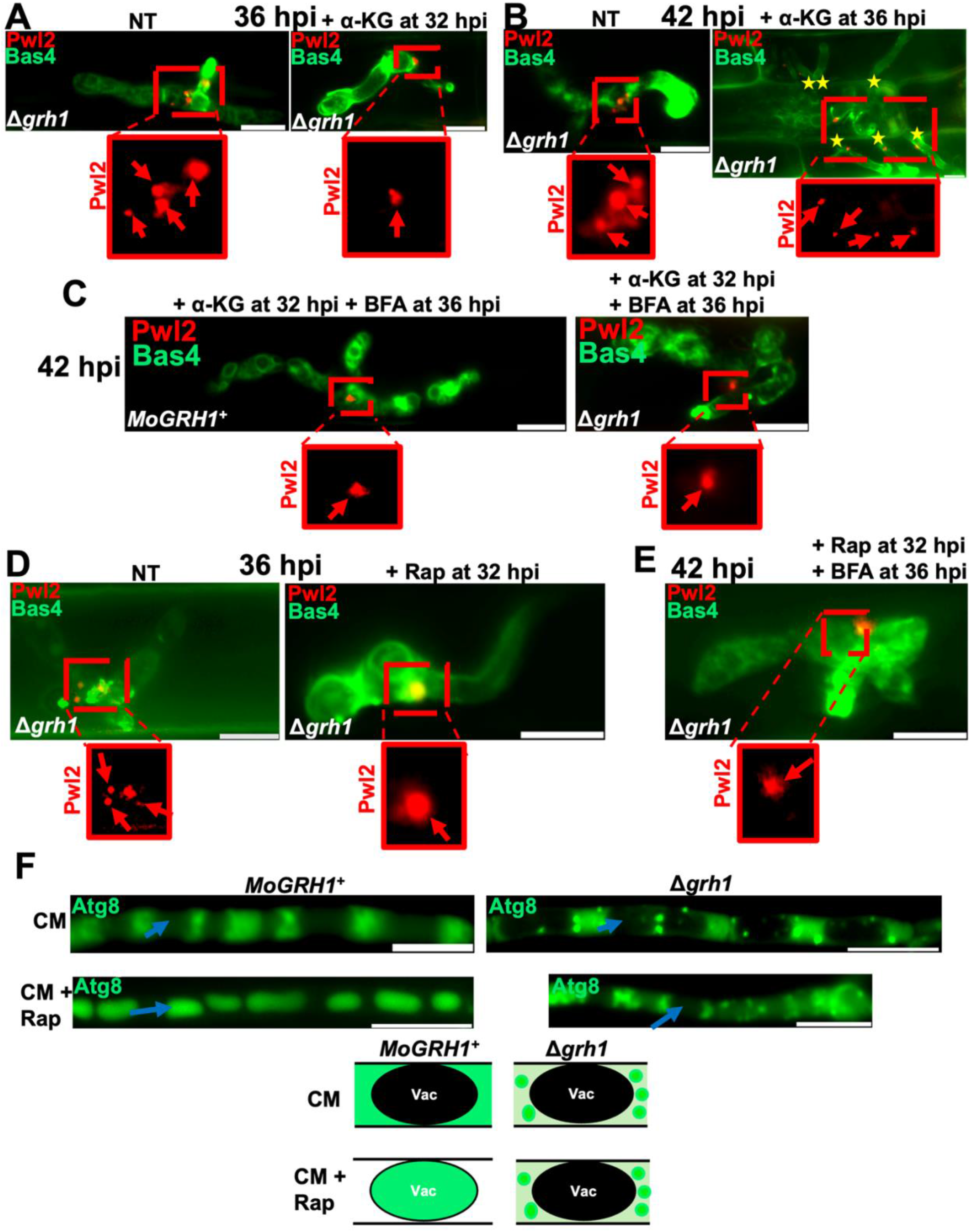
UPS in Δ*grh1* is restored by α-ketoglutarate or rapamycin treatment. **A-C.** Live-cell imaging of detached rice leaf-sheath epidermal cells at 36 hpi (**A**) and 42 hpi (**B,C**) infected with the indicates strains. Samples were either untreated (NT), treated with 10 mM α-ketoglutarate (DMKG) at 32 hpi (**A**) or 36 hpi (**B**), or treated with DMKG at 32 hpi and with 50 µg ml^-1^ BFA in 0.1 % DMSO at 36 hpi (**C**). **D,E.** Live-cell imaging of the indicated strains in detached rice leaf-sheath epidermal cells at 36 hpi (**D**) and 42 hpi (**E**). Samples were either untreated (NT), treated with 10 µM rapamycin (Rap) at 32 hpi (**D**), or treated with 10 µM rapamycin (Rap) at 32 hpi and with 50 µg ml^-1^ BFA in 0.1 % DMSO at 36 hpi (**E**). **A-E.** Red arrows indicate BICs or BIC fragment for clarity. Yellow stars mark IH extending into neighboring host cells. Representative images were selected from 50 infected rice cells examined per leaf sheath per treatment across three biological replicates. Scale bars =10 μm. **F.** The indicated strains expressing *GFP-ATG8* were cultured in liquid CM for 42 hr. Mycelia were harvested, washed with water, and transferred to fresh CM alone or CM supplemented with 10 µM rapamycin (Rap), as indicated, for 4 hr before imaging. Blue arrows indicate examples of vacuoles. The schematic summarizes the treatment outcomes; vac denotes vacuole. Images were selected from observing 50 mycelial fragments, repeated in triplicate. Scale bars = 10 μm.

Removing the need for autophagy restored UPS cargo sorting and pathway integrity in Δ*grh1* mutant strains treated with DMKG, prompting us to ask whether constitutively activating autophagy may have a similar remediating effect. In yeast, Grh1 is required for secretory autophagosome biogenesis, which is not induced in response to the TOR pathway inhibitor rapamycin, but Grh1 is not required for canonical degradative autophagy and the formation and trafficking of degradative autophagosomes, which is activated by rapamycin (Bruns et al. 2011). We treated *MoGRH1^+^* and Δ*grh1* infected rice leaf sheaths with rapamycin, finding that in *MoGRH1^+^*, rapamycin did not affect UPS or conventional effector secretion, but it did curtail IH growth (**Supplementary Fig. S4A,B**), as previously reported for *M. oryzae* WT strains (Sun et al. 2018; Li et al. 2023a). In the Δ*grh1* strain, rapamycin treatment restored single intact BICs in each infected rice cell (**Fig. 6D**), which dual rapamycin and BFA treatments showed coincided with restoration of the UPS pathway (**Fig 5E**). Therefore, constitutively activating degradative autophagy in IH with rapamycin restored cargo sorting and vesicle trafficking via UPS in Δ*grh1*, but inhibited (*MoGRH1^+^*) or failed to remediate (Δ*grh1*) IH growth. In vegetative hyphae, in the *MoGRH1^+^* strain, rapamycin induced degradative autophagy in CM, resulting in the trafficking of autophagosomes to vacuoles and the accumulation of GFP fluorescence in end-stage autophagy, but rapamycin did not produce such autophagic flux in Δ*grh1* hyphae, where GFP-Atg8-labelled puncta were observed in cytoplasm but vacuoles remained empty of the GFP signal (**Fig. 6F**). Thus, constitutively activating degradative autophagy with rapamycin, while not remediating end-stage degradative autophagy in Δ*grh1*, was nonetheless sufficient to restore UPS and cargo sorting in the mutant strain. These results are consistent with a role for MoGrh1, like the GRASP55 orthologue, in enhancing autophagosome-vacuole/lysosome fusion (Zhang et al. 2018), and also support the notion that rapamycin-induced degradative autophagosomes that do not fuse to the vacuole in Δ*grh1* can be repurposed as secretory autophagosomes to remediate UPS, such as is observed when degradative autophagosome-lysosome/ vacuole fusion is blocked in nerve cells leading to increased secretory autophagosome production (Ejlerskov et al. 2013). How these compartments are repurposed as secretory autophagosomes remains to be determined. We conclude that MoGrh1 plays a critical role in balancing IH growth, degradative and secretory autophagy, and the sorting and organizing of effector vesicular trafficking during living host plant cell colonization.

## Discussion

Fungal plant pathogens threaten global food security. One of the most notorious is the blast fungus *M. oryzae*, which infects a wide range of grasses including rice and wheat (Wilson and Talbot, 2009; Wilson, 2021; Valent, 2025). Early infection involves a biotrophic growth phase, where fungal IH colonize successive living host rice cells (Kankanala et al. 2007). Understanding more about effector secretion, which occurs during biotrophy via two distinct pathways, would fill gaps in our knowledge of effector biology and aid the development of more robust strategies for tackling blast and other plant diseases. However, although previous work demonstrated a requirement for the SNARE MoSso1 and the exocyst complex in mediating *M. oryzae* UPS (Giraldo et al. 2013), how effectors are sorted and trafficked through the different pathways, how the pathways are organized, and what other components and cellular relationships are involved, is unknown. Here, we showed how the *M. oryzae* orthologue of the yeast ERGIC protein Grh1, MoGrh1, is required for sorting cytoplasmic effectors into the BFA-insensitive UPS pathway under the native autophagy-inducing conditions of host cell infection. In the absence of MoGrh1, UPS ceased, and cargo was instead trafficked to multiple BIC-like surface puncta (and intracellular patches) in a disorganized manner via a BFA-sensitive pathway. One significant outcome of our findings is the unique opportunity the *M. oryzae* Δ*grh1* strain affords us to understand what happens when cytoplasmic effectors are trafficked via a non-physiological secretion pathway. Although some Pwl2 accumulated internally, Pwl2 still deployed in the Δ*grh1* mutant strain to external (albeit multiple) subapical BIC-like structures, indicating that cytoplasmic effectors carry currently unknown motifs or intrinsic features (Dulal and Wilson, 2024) signposting them to external BIC structures regardless of the secretion pathway used. However, the formation of Δ*grh1* split BICs, which became single, intact BICs when UPS was restored by α-ketoglutarate or rapamycin treatment, tells us that UPS provides directional, focused and controlled secretion into the BIC that is unmatched by conventional ER-Golgi secretion but essential for maintaining subapical BIC integrity. This may provide a physiological basis (targeted, focal subapical secretion) for UPS function and explain why *M. oryzae* has evolved to use UPS to deploy plant cell-destined cytoplasmic effectors into the BIC.

### Role of the ERGIC in effector secretion

By using MoGrh1-GFP as an *M. oryzae* ERGIC marker, and by finding that Pwl2-mCherry:HDEL is a transient ERGIC cargo that co-localizes with MoGrh1 when trapped by BFA, we illuminated the first BFA stable ERGIC-like remnants (Marie et al. 2008; Saraste and Prydz, 2026) in filamentous fungi. Yeast and mammalian ERGIC structures are sorting organelles for both vesicular UPS and conventional ER-Golgi secretion cargo with connections to recycling endosomes and (in animals) the Golgi stack (Saraste and Prydz, 2026), as well as to ER exit sites (Bruns et al. 2011; Tojima et al. 2024; Tojima and Nakano, 2026). Autophagosome membrane precursor production occurs through the formation of ERGIC-derived COPII vesicles (Ge et al. 2017). Our work implicates BFA-resistant ERGIC-like structures in *M. oryzae* as MoGrh1-dependent central sorting hubs for cytoplasmic effectors, warranting their further interrogation.

### Role of MoGrh1 in autophagy

MoGrh1 has a role in Type IV UPS and degradative autophagy, which contrasts with observations in yeast but is in line with known roles of mammalian GRASPs. In yeast, Grh1 is required for secretory autophagosome biogenesis from specialized compartments formed under starvation conditions (Duran et al. 2010; Bruns et al. 2011), potentially from interactions between the ERGIC and endosomes (Tojima et al. 2024; Saraste and Prydz, 2026). Yeast secretory autophagosomes avoid fusion with the vacuole and subsequent cargo degradation, and delivery to the PM requires endosomal sorting (Duran et al. 2010). Yeast Grh1 is not required, however, for degradative autophagosome biogenesis, which unlike secretory autophagosome biogenesis is induced by rapamycin (Duran et al. 2010; Bruns et al. 2011). Furthermore, yeast Grh1 mediates the secretory autophagosome biogenesis required for Type III UPS of leaderless Acb1 (Duran et al. 2010; Bruns et al. 2011), but it has not yet been shown to be involved in yeast Type IV UPS (Saraste and Prydz, 2026).

In contrast to yeast Grh1, animal GRASP orthologues are required for both Type III and Type IV UPS (Saraste and Prydz, 2026), in addition to degradative autophagy (Zhang et al. 2018). Mammalian GRASP55 (but not GRASP65) localizes to secretory autophagosomes and multivesicular bodies under cellular stress conditions, including rapamycin treatment, to induce UPS (Nuchel et al. 2021). GRASP55 may facilitate secretory autophagosome fusion to the PM (Zhang et al. 2018), and it also functions as a membrane linker to enhance autophagosome-lysosome/vacuole fusion in end-stage degradative autophagy (Zhang et al. 2018). GRASP55 is required for the unconventional secretion of both leaderless and signal peptide-carrying cargo (i.e. Type III and Type IV UPS) involving at least 63 proteins evenly split between the two classes (Nuchel et al. 2020). mTORC1 inhibition by rapamycin in mammalian cells induced both degradative autophagy and secretory autophagy simultaneously (although the composition of the different classes of autophagosomes was different) (Nuchel et al. 2021). Here, our results show MoGrh1 functions similarly to GRASP55, firstly by being involved in Type IV UPS, and secondly, by being required for degradative autophagy, likely at the autophagosome-vacuole fusion level. Consistent with a role in autophagosome-vacuole fusion, MoGrh1-GFP localized around vacuoles during autophagy activation (but formed puncta during autophagic flux minima in IH at 40 hpi and at later timepoints following α-ketoglutarate treatment), while GFP-Atg8-labelled vesicles accumulated in the Δ*grh1* cytoplasm.

The loss of *MoGRH1* reduced or mis-timed end-stage degradative autophagy under both axenic shake and biotrophic growth conditions. Rapamycin treatment restored UPS in Δ*grh1*, but end-stage degradative autophagy remained impaired. In nerve cells, increasing the pool of degradative autophagosomes, e.g. by blocking degradative autophagosome-lysosome fusion with Bafilomycin A (BafA1), switched degradative autophagosomes to secretory autophagosomes available for UPS (Ejlerskov et al. 2013), suggesting that degradative and secretory autophagy are inextricably linked (Sawai et al. 2024). Our results support a similar scenario such that treatment with rapamycin that constitutively activates autophagy would be expected to increase degradative autophagosome production in Δ*grh1* but, because they cannot fuse efficiently to vacuoles, the pool of degradative autophagosomes available for switching to secretory autophagosomes would also increase, potentially explaining how rapamycin restores UPS but not end-stage degradative autophagy in Δ*grh1* IH. Commensurate with this notion, and consistent with end-stage autophagy being unnecessary for secretory autophagy in yeast (Manjithaya and Subramani, 2011), BafA1 treatment, which inhibits autophagosome-lysosome/vacuole fusion and thus end-stage autophagy, did not affect conventional or unconventional secretion in *M. oryzae* (Sun et al. 2018). Additional evidence for the involvement of secretory autophagosomes in *M. oryzae* UPS comes from previous work showing UPS requires the t-SNARE Sso1 ortholog (Giraldo et al. 2013), a required receptor for secretory autophagosome membrane fusion but not for degradative autophagosome membrane fusion (Duran et al. 2010). When also considering that rapamycin-induced degradative autophagy inhibited WT IH growth, our results suggest MoGrh1 mediates degradative and secretory autophagy homeostasis during biotrophy to coordinate effector secretion with IH growth needs.

Despite the importance of *MoGrh1*-dependent autophagy to UPS, removing the need for autophagy by treating infected rice cells with DMKG restored UPS in Δ*grh1* while having no effect on secretion in *MoGRH1^+^*, suggesting that under these artificial nutrient-replete conditions, autophagy balance regulated by MoGrh1 is not required, and the UPS pathway remains active and potentially supported by other vesicular processes. Different types of UPS utilize different vesicles (Nuchel et al. 2021), and it could be that DMKG treatment alters ERGIC interactions, connecting UPS sorting (which no longer requires MoGrh1) to other cellular processes that remain to be determined but that are no longer limited by nutrient stress. Tellingly, the maintenance of UPS under nutrient-replete conditions in the absence of MoGrh1 shows that, like in neurons (Saraste and Prydz, 2026), UPS is constitutive in *M. oryzae* during biotrophy and is not simply, as in yeast, a response to nutrient stress. This is potentially more like the situation in animal cells, where UPS responds to a range of cellular signals and stresses (Nuchel et al. 2021). In *M. oryzae*, effector genes are only expressed *in planta*, and although a global regulator is currently unknown, it would make sense for UPS to be activated concurrently with effector gene expression regardless of cellular or environmental stress cues, potentially by a MAP kinase such as Pmk1 (Wilson and Talbot, 2009) responding to multiple external and developmental stimuli. Taken together, we conclude that cytoplasmic effector secretion via UPS is constitutive and may be linked to infection-related development, but it is critically supported by MoGrh1-dependent autophagy under the native nutrient-restrictive conditions of biotrophic growth.

### Role of MoGrh1 in secretion

Yeast Grh1 secretion function has focused on Type III UPS of Acb1 (Bruns et al. 2011), but a comprehensive analysis of GRASP55-dependent UPS cargo in human cell lines revealed its role in both Type III and Type IV UPS under stress conditions including nutrient stress (Nuchel et al. 2021; Ahat et al. 2022). The loss of GRASP55 abolished UPS cargo secretion by any route (Nuchel et al. 2021), which is in contrast to our results where secretion was misrouted rather than abolished in Δ*grh1*. This suggests that the ER-Golgi pathway is the default secretion pathway in *M. oryzae* (at least for SP-carrying cargo), and MoGrh1 is required (under autophagy-inducing conditions) for actively sort cytoplasmic effectors into the UPS pathway. Because cytoplasmic effectors carry no known motifs distinguishing them from apoplastic effectors (Dulal and Wilson, 2024), future work may comprehensively catalogue these important mediators of plant-microbe interactions by, for example, comparing the Δ*grh1* and WT secretome, such as was performed for GRASP55 in human cell lines (Nuchel et al. 2021). Such secretome analysis may also determine if MoGrh1 is required for Type III UPS, which is undescribed in *M. oryzae*. Finally, it is conceivable that UPS cargo is ultimately delivered to external BICs in exosomes (Richter et al. 2024); future work focused on understanding precisely how MoGrh1-dependent UPS interacts with the BIC to maintain its integrity may shed light on the post-PM stage of the cytoplasmic effector delivery process.

## Conclusion

We conclude that UPS in *M. oryzae* during biotrophy is constitutive but critically supported by MoGrh1-dependent secretory autophagy and effector sorting under the nutrient-stress conditions of living host rice cell colonization. This work highlights ERGIC-like structures as hubs for cytoplasmic effector trafficking and provides new insights into unconventional secretion that may be applicable to a wide range cell types as well as to pathogens with similar infection strategies.

## Materials and Methods

### Fungal strains, culture conditions and physiological analyses

The *M. oryzae* isolate Guy11 was used as the parental wild-type (WT) strain. All fungal strains used in this study are listed in **Supplementary Table S1**. Strains were cultured on solid complete media (CM; Li et al. 2015) or 3 % water agar media (Li et al. 2023a), or in liquid CM, at 26 °C. For plate-based assays, cultures were maintained at 26 °C under a 12 hr light/12 hr dark cycle for 10–14 days. Conidia were harvested from 10-day-old cultures on CM plates, counted using a haemocytometer, and used for the indicated assays, unless otherwise stated. Each experiment was performed with at least three independent biological replicates.

For vegetative hyphal treatments, mycelia were cultured in liquid CM for 42 h, harvested, washed three times with distilled water, and transferred to the indicated treatment medium for up to 4 hr before imaging or further analysis.

### Vector construction

*M. oryzae* gene sequences were obtained from the *M. oryzae* genome database (http://fungi.ensembl.org/Magnaporthe_oryzae/Info/Index) and used for vector construction. To delete the *MoGRH1* coding sequence at locus MGG_05757, we first constructed a vector, GRASPKO:pNEB, carrying the sulfonylurea-resistance marker *ILV2* flanked by 1kb of up- and downstream *MoGRH1* sequences. The *MoGRH1* flanks and *ILV2* coding sequence were amplified by PCR and subcloned into pNEB using the yeast gap repair method. Primers were designed to introduce terminal sequences homologous to the neighboring assembly fragments, thereby arranging the deletion cassette in the order 5′ *MoGRH1* 1 kb flank–*ILV2*–3′ *MoGRH1* 1 kb flank. The terminal fragments also contained homology to the ends of the linearized pNEB yeast-recombination vector. Linear PCR fragments were amplified with Phusion High-Fidelity DNA Polymerase (Thermo Fisher Scientific). Oligonucleotides were designed using PrimerQuest (Integrated DNA Technologies). *MoGRH1* flanking regions were amplified from Guy11 DNA that had been extracted using our typical method (Li et al. 2015). The *ILV2* coding region was amplified from pCB1532 (Wilson et al. 2010). The three purified amplicons and linearized pNEB vector, which carries the *URA3* gene, were co-transformed into a *ura3* auxotrophic *Saccharomyces cerevisiae* strain using the Yeast Transformation Kit (Sigma Aldrich, USA), and transformants were selected on synthetic complete medium lacking uracil. Individual Ura+ colonies were cultured on selective medium lacking uracil, and recombinant plasmid DNA was recovered using the Zymoprep Yeast Plasmid Miniprep II kit (Zymo Research, USA). The kit uses Zymolyase-mediated cell-wall digestion followed by alkaline lysis and spin-column purification. The recovered plasmids were transformed into competent *E. coli* cells (DH5α), selected by ampicillin resistance conferred by the *ampR* gene on pNEB, and purified from individual colonies. Correct assembly of GRASPKO:pNEB was confirmed by PCR.

The constructs RP27-*GRASP*-GFP-pBV591 and Pwl2-mCherry-HDEL; RP27-*GRASP*-GFP-pBV591 were generated by GenScript USA, Inc., using pBV591 (Khang et al. 2010) as the backbone. The indicated sequences were synthesized, cloned into the respective vectors using the CloneEZ cloning method, and verified by sequencing.

### Generation of mutant strains

To investigate the function of *MoGRH1* in *M. oryzae*, the first 1 kb of the *MoGRH1* coding sequence was replaced with the sulfonylurea-resistance marker *ILV2* by homologous recombination using the PCR-based split-marker method (Wilson et al. 2010). Using the GRASPKO:pNEB construct described above as the template, we amplified 1kb of each flanking region fused to an overlapping region of *ILV2*. WT Guy11 protoplasts were prepared as previously described (Li et al. 2015), transformed with the amplified DNA fragments, and transformants were selected on plates for resistance to sulphonyl urea (Chlorimuron ethyl, Thermo Fisher Scientific, USA). Candidate deletion mutants were identified by PCR using the primers listed in **Supplementary Table S2**. Five independent Δ*grh1* mutants with indistinguishable phenotypes were confirmed by initial characterization of growth, sporulation and infection rates.

To generate the Δ*grh1* strain expressing *PWL2-mCherry:NLS* and *BAS4-GFP*, we deleted *MoGRH1* (as described above) in our previously generated Guy11 strain (Li et al. 2023a) carrying the pBV591 vector (Khang et al. 2010) expressing these genes.

To generate the Δ*grh1* strain expressing *GFP-ATG8*, we transformed protoplasts of Δ*grh1* with our previous vector GFP-ATG8_pGTN (Li et al. 2023a) and selected transformants based on geneticin resistance and confirmation by PCR using the primers in **Supplementary Table S2**.

To generate strains expressing either *MoGRH1-GFP* alone or *PWL2-mCherry-HDEL* and *MoGRH1-GFP*, protoplasts were transformed with the RP27-*GRASP*-GFP-pBV591 or Pwl2-mCherry-HDEL; RP27-*GRASP*-GFP-pBV591 vectors described above. Transformants were selected based on Hygromycin B resistance (RPI Research Products, USA) and verified by PCR using the primers listed in **Supplementary Table S2**.

For each vector, PCR-confirmed transformants were evaluated by fluorescence microscopy in detached rice leaf sheaths. Because vectors carrying genes of interest integrate randomly into the *M. oryzae* genome, at least three independent transformants for each construct in each strain background were confirmed in detached rice leaf sheath assays as being equivalent with regards to labelled-protein production levels and localization. One representative transformant per construct and strain background was used for subsequent analyses.

### Plant inoculation and infection assays

For whole-plant inoculation assays, conidia were collected from 14-day-old cultures grown on solid CM and suspended in 0.2 % gelatin at 1×10^5^ conidia ml^-1^. Spore suspensions were sprayed onto 3-week-old seedlings of the susceptible rice cultivar CO-39. Infected leaves were harvested, dried, and photographed at 5 days post-inoculation (dpi). Images shown are representative of at least five leaves from each of three independently inoculated plants per strain.

Appressorium formation was assessed on artificial hydrophobic surfaces. A conidial suspension at conidia 1×10^4^ ml^-1^ was applied to the plastic coverslips surface, and appressorium formation was quantified at 24 hr post-inoculation (hpi) by scoring 50 conidia per replicate. Appressorial penetration was evaluated in detached rice leaf sheaths by determining the proportion of 50 appressoria that penetrated underlying epidermal cells at 30 hpi. Cell-to-cell movement of invasive hyphae (IH) was quantified as the proportion of 50 initially infected rice leaf-sheath epidermal cells containing IH in adjacent host cells at 44 hpi. Each assay was repeated in three independent biological replicates.

### Live-cell imaging and chemical treatments

Detached rice leaf sheaths were prepared and inoculated with 0.2 % gelatin containing 0.5 ×10^5^ conidia ml^-1^ of the indicated fungal strains. Images were obtained at the specified time points using a Nikon Eclipse Ni-E upright microscope with NIS-Elements software. GFP and mCherry were imaged using excitation/emission settings of 488 nm/505–531 nm and 543 nm/590–632 nm, respectively. Three-dimensional reconstructions were generated using the volume-rendering function in Nikon NIS-Elements BR software.

Brefeldin A (BFA), rapamycin (Rap), and dimethyl α-ketoglutarate (DMKG) were obtained from Sigma-Aldrich (USA). For BFA assays, infected detached rice leaf sheaths were treated at 32 hpi with 50 µg ml^-1^ BFA in 0.1 % DMSO and imaged at 36 hpi, unless otherwise stated. Control samples received 0.1% DMSO alone. For rapamycin treatment, infected leaf sheaths were treated with 10 µM Rap (from a 25 mg ml^-1^ stock in 0.1 % DMSO) at 32 hpi and imaged at 36 hpi, unless otherwise stated. Infected leaf sheaths were treated with 10 mM DMKG (from a 30 mg/ml stock in 0.1 % DMSO) at 32 hpi and imaged at 36 hpi, unless otherwise stated. For sequential DMKG and BFA treatments, samples were treated with 10 mM DMKG at 32 hpi, followed by 50 µg ml^-1^ BFA at 36 hpi, and imaged at 42 hpi. For sequential Rap and BFA treatments, samples were treated with 10 µM Rap at 32 hpi, followed by 50 µg ml^-1^ BFA at 36 hpi, and imaged at 42 hpi. All chemical-treatment experiments were performed independently on at least three different leaf sheaths, with representative images from examining 50 infected rice cells per leaf sheath shown.

For vegetative mycelia, 10 mM DMKG and 10 µM Rap were added to liquid shakes, and images were taken after 4 hrs. Images are representative of three independent shake cultures per treatment.

For autophagic flux analysis, rates were calculated by observing strains *in planta* at the indicated time points, with and without DMKG treatment, and scoring the percentage of vacuoles exhibiting at least some GFP fluorescence in 50 infected rice cells, repeated in triplicate.

### Statistical analysis

Rates of sporulation, appressorium formation, appressorial penetration, IH cell-to-cell movement and autophagic flux were analyzed using R version 4.2.2. Mean ± SD values among strains and treatments were compared using the Agricolae package (version 1.3-7), and graphs were generated using ggplot2 (version 3.5.1). Statistical significance was assessed by one-way analysis of variance (ANOVA) followed by Tukey’s honestly significant difference (HSD) test. Differences were considered significant at *P* ≤ 0.05.

## Acknowledgements

This work was supported by National Science Foundation funding to R.A.W. (PBI IOS-2512144).

## Author contributions

R.A.W. conceived and supervised the project and acquired funding. N.D. performed the experiments. N.D. and R.A.W. analyzed the data. R.A.W. wrote the manuscript. N.D. reviewed and edited the manuscript.

---

**Supplementary Figure S1.**
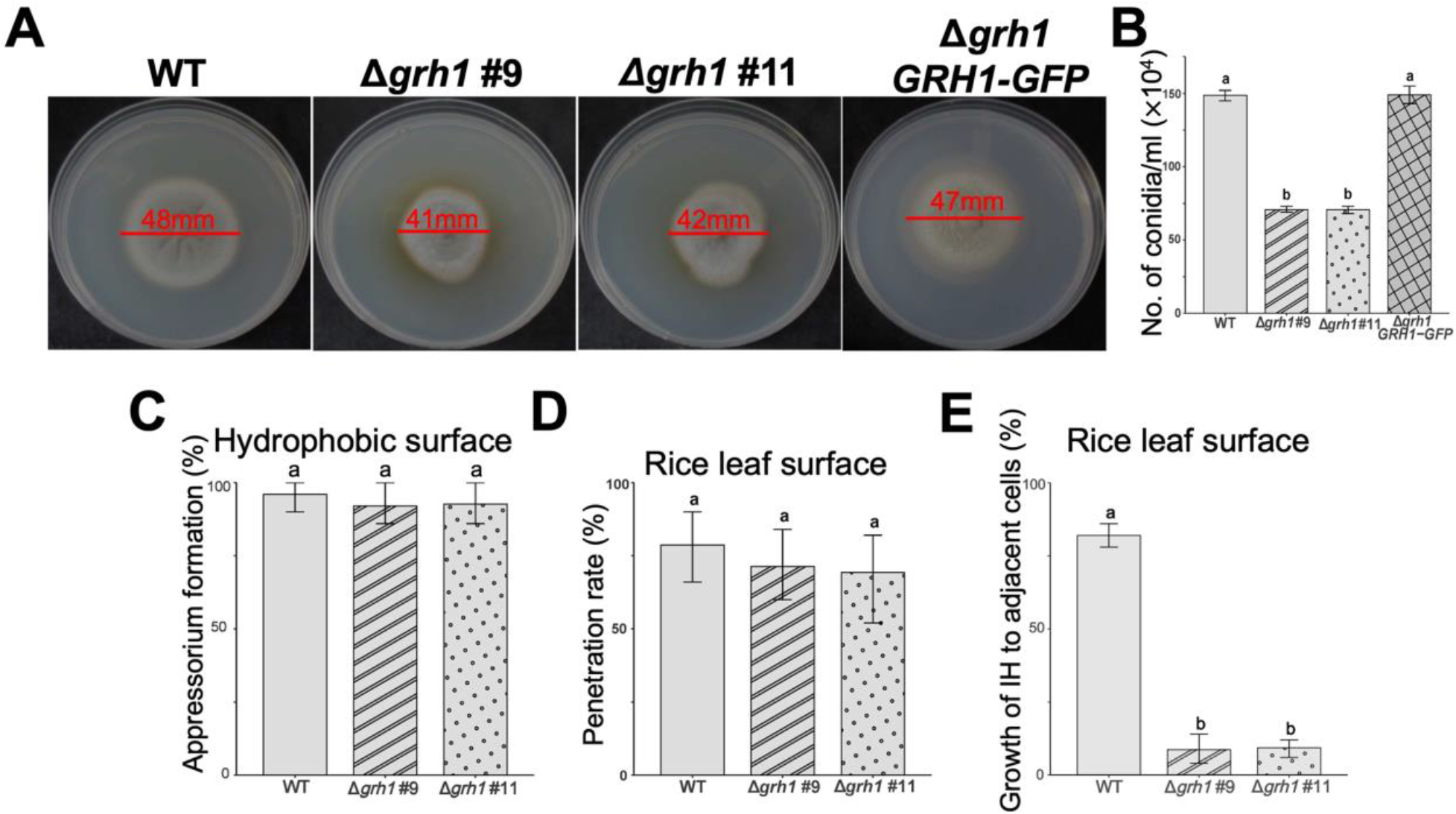
Loss of *MoGRH1* affects sporulation and biotrophic growth. **A.** Growth tests on complete media (CM) of the indicated strains after 10 days post inoculation. Representative images are shown from three biological replications. Values are the average diameters from three plates per strain. **B.** Sporulation rates of the indicated strains. Spores were harvested from CM at 10 days, and the number of spores ml^-1^ (×10^4^) were counted, repeated in triplicate. **C.** Appressorium formation rates of the indicated strains by 24 hpi. **D.** Appressorium penetration rates of the indicated strains by 32 hpi, observed by live-cell imaging of infected detached rice leaf sheaths. **E.** Cell-to-cell movement rates of the indicated strains by 44 hpi, observed by live-cell imaging of infected detached rice leaf sheaths. IH movement was scored if at least one new strand of invasive hyphae (IH) was present in adjacent cells. **B-E.** Values are means ± SD. Rates were compared among strains using one-way ANOVA followed by Tukey’s HSD test. For each graph, different letters above columns indicate significant differences of the means at *P* ≤ 0.05. **C-E**. Values were obtained from observing 50 germinating spores on artificial hydrophobic surfaces (plastic coverslips) (**C**), 50 appressoria on the surface of each detached rice leaf sheath (**D**) and 50 penetration sites in detached rice leaf sheath cells (**E**), repeated in triplicate.

**Supplementary Figure S2.**
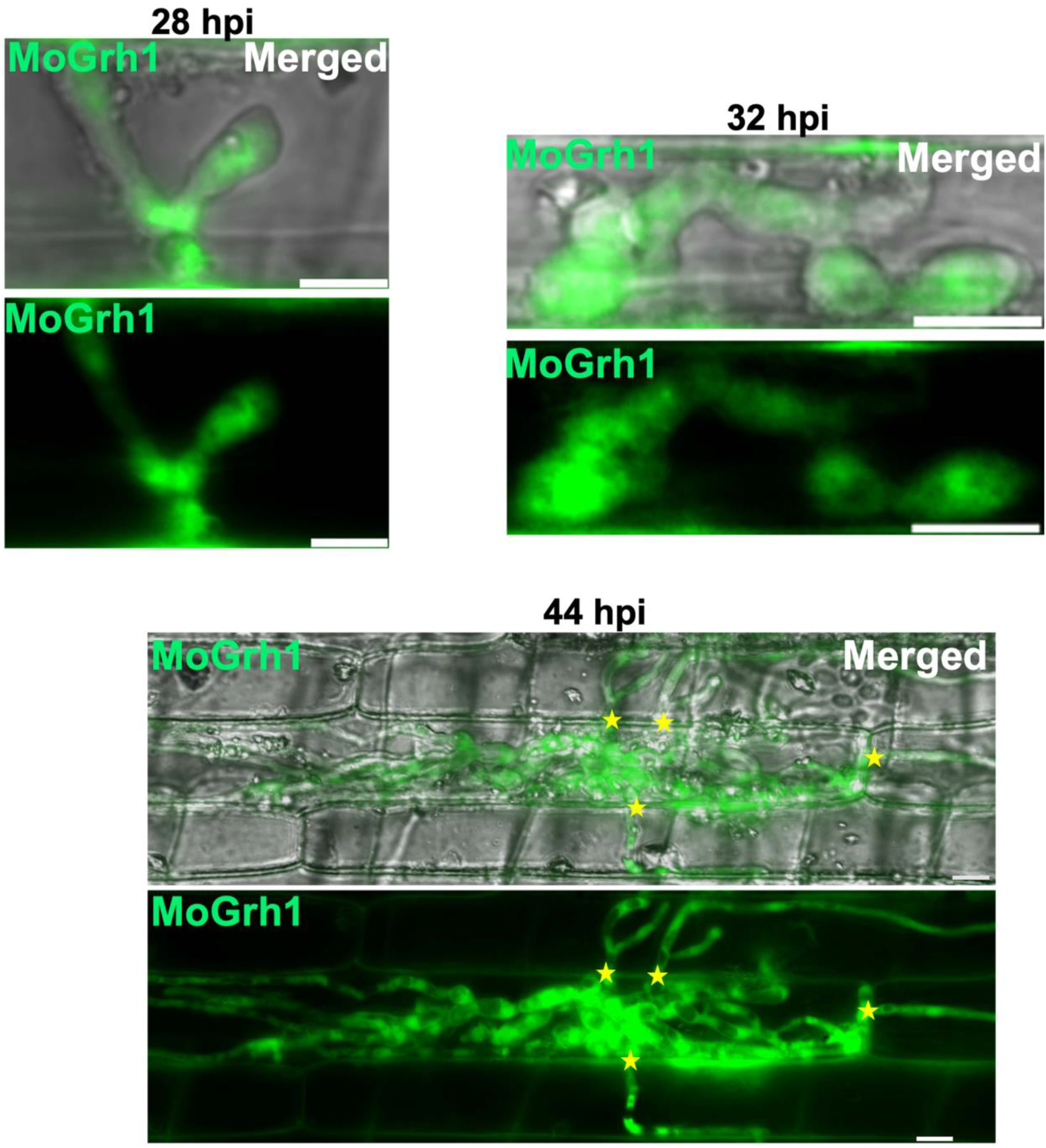
MoGrh1 localization in IH in rice cells is dynamic as biotrophy progresses. Live-cell imaging of the Δ*grh1 MoGRH1-GFP* complementation strain in detached rice leaf-sheath epidermal cells at the indicated time points. Yellow stars mark IH from penetration sites extending into neighboring host cells. Representative images were selected from observations of 50 infected rice cells per leaf sheath per timepoint, with experiments performed in triplicate. Scale bars = 10 μm.

**Supplementary Figure S3.**
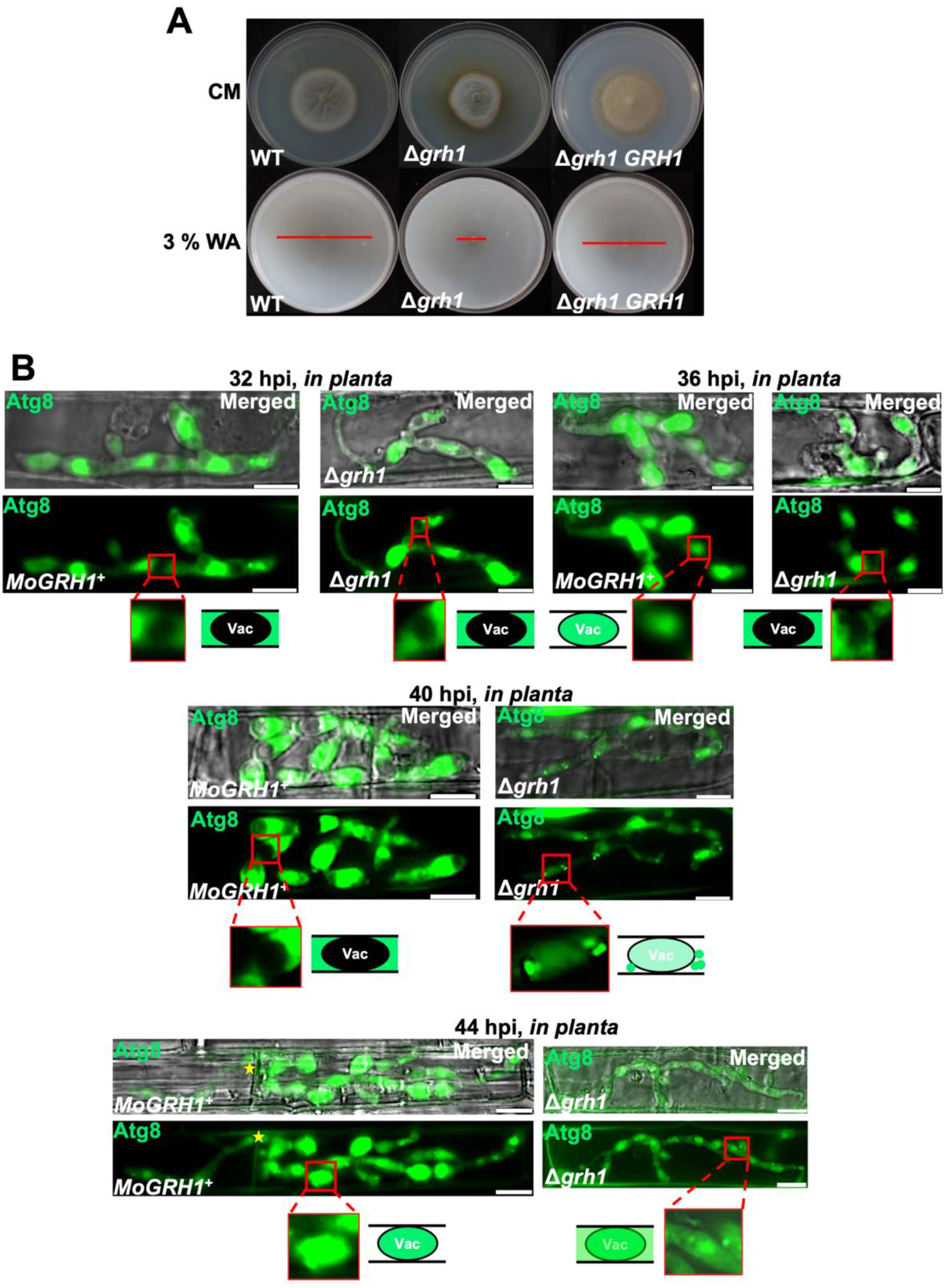
Loss of *MoGRH1* impairs autophagy on plates and *in planta*. **A.** Plate growth tests of the indicated strains on complete media (CM) and nutrient-free 3% water agar (WA). Red bars indicate colony diameters of starvation mycelia growing from the central plug. Images were captured at 10 days post inoculation. **B.** Live-cell imaging of detached rice leaf-sheath epidermal cells infected with the indicated strains expressing *GFP-ATG8* at the indicated time points. Vac denotes vacuole. Stars indicate IH extending into neighboring host cells. Representative images were selected from 50 infected rice cells examined per leaf sheath for each treatment across three biological replicates. Scale bars = 10 μm.

**Supplementary Figure S4.**
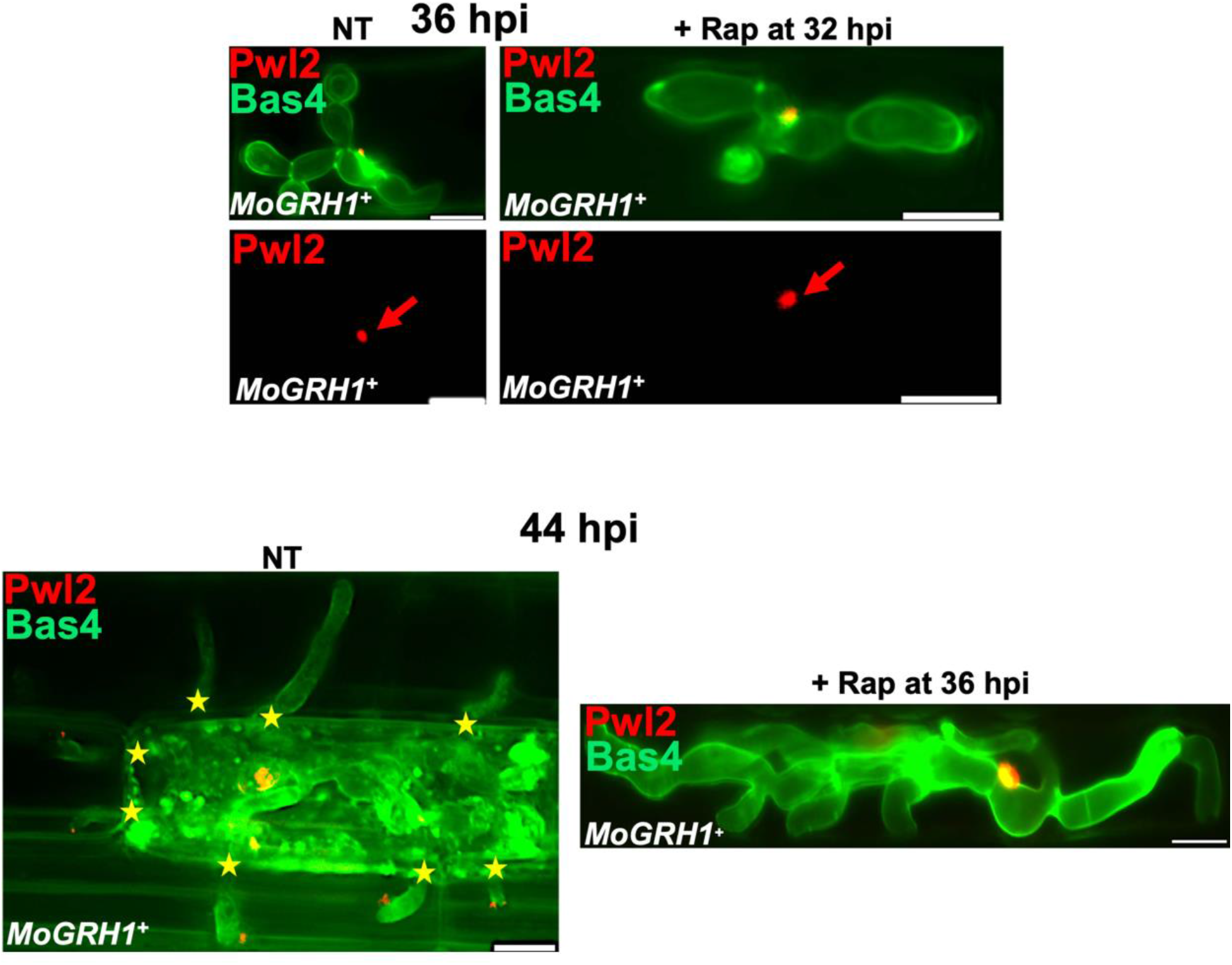
Rapamycin treatment inhibits *M. oryzae* IH growth and branching but not effector secretion. **A, B.** Live-cell imaging of detached rice leaf-sheath epidermal cells at 36 hpi (**A**) and 44 hpi (**B**) infected with the indicated strain. Samples were either untreated (NT) or treated with 10 µM rapamycin (Rap) at 32 hpi and viewed at 36 hpi (**A**) or treated at 36 hpi and viewed at 44 hpi (**B**). Red arrows indicate BICs for clarity. Yellow stars mark IH extending into neighboring host cells. Representative images were selected from 50 infected rice cells examined per leaf sheath for each treatment across three biological replicates. Scale bars = 10 μm.

