## Supplementary Table S1 for "Blast fungus unconventional effector secretion sorting requires pre-Golgi compartment protein MoGrh1"

**Supplementary Table S1: Strains used**

| **Genotype** | **Description** | **Source** |
| --- | --- | --- |
| WT | *M. oryzae* wild type Guy11 isolate used in this study. | Wilson et al. 2010 |
| Δ*grh1* | Guy11 parental strain carrying a deletion of the *MoGRH1* gene (MGG_05757) encoding the *M. oryzae* GRASP55 orthologue. | This study |
| *MoGRH1^+^ PWL2-mCherry:NLS; BAS4-eGFP* | Guy11-derived strain expressing *PWL2-mCherry:NLS* and *BAS4-eGFP* from native promoters following genomic integration of the pBV591 vector (Khang et al. 2010). | Li et al. 2023a |
| Δ*grh1* *PWL2-mCherry:NLS; BAS4-eGFP* | Δ*grh1* strain expressing *PWL2-mCherry:NLS* and *BAS4-eGFP* from native promoters, derived from the deletion of *MoGRH1* in the *PWL2-mCherry:NLS; BAS4-eGFP* expressing strain. | This study |
| Δ*grh1* *MoGRH1-eGFP* | Δ*grh1* parental strain complemented with *MoGRH1-GFP* after genomic integration of a modified pBV591 plasmid expressing *MoGRH1* under the *RP27* promoter in place of *BAS4* and its promoter. | This study |
| *PWL2-mCherry:HDEL; PrRP27:MoGRH1-eGFP* | Guy11-derived strain expressing, under the *PWL2* promoter, *PWL2* fused to *mCherry* but with the *NLS* replaced by four codons encoding the ER retention signal HDEL, also expressing *MoGRH1* fused to *eGFP* under the *RP27* promoter, after genomic integration of a modified pBV591vector. | This study |
| *MoGRH1^+^* *GFP-ATG8* | Guy11-derived strain expressing *GFP-ATG8* after transformation with the pGTN vector carrying *GFP-ATG8*. | Li et al. 2023a |
| Δ*grh1* *GFP-ATG8* | Δ*grh1*-derived strain expressing *GFP-ATG8* after transformation with the pGTN vector carrying *GFP-ATG8*. | This study |

Wilson RA, Gibson RP, Quispe CF, Littlechild JA, Talbot NJ (2010) An NADPH-dependent genetic switch regulates plant infection by the rice blast fungus. Proc Natl Acad Sci USA 107: 21902-21907. doi: 10.1073/pnas.1006839107

Khang CH, Berruyer R, Giraldo MC, Kankanala P, Park SY, Czymmek K, Kang S, Valent B (2010) Translocation of *Magnaporthe oryzae* effectors into rice cells and their subsequent cell-to-cell movement. Plant Cell 22: 1388-1403. doi: 10.1105/tpc.109.069666

Li G, Gong Z, Dulal N, Marroquin-Guzman M, Rocha RO, Richter M, Wilson RA (2023a) A protein kinase coordinates cycles of autophagy and glutaminolysis in invasive hyphae of the fungus *Magnaporthe oryzae* within rice cells. Nat Commun 14: 4146. doi: 10.1038/s41467-023-39880-w
