## Supplementary Table S2 for "Blast fungus unconventional effector secretion sorting requires pre-Golgi compartment protein MoGrh1"

| **Primer Name** | **Sequence** | **Use** |
| --- | --- | --- |
| GRASPKO-LF-Fw | AACTGTTGGGAAGGGCGATCGGTGCGGGCCATGACGCGGTCAAAGGA | Amplification of *MoGRH1* flanks for gene deletion |
| GRASPKO-LF-Re | TGTGGGGCACTGTGGCGTTGGCACGTCGACATGGGCGTTGTTATCCTTCTG | Amplification of *MoGRH1* flanks for gene deletion |
| GRASPKO-RF-Fw | TGCACGGGAATTGCATGCTCTCACGTCGACGTAGAGCGTTGTAGAGCATAGAC | Amplification of *MoGRH1* flanks for gene deletion |
| GRASPKO-RF-Rw | TTCACACAGGAAACAGCTATGACCATGATTCCGCGACTCGAATCAAAGATA | Amplification of *MoGRH1* flanks for gene deletion |
| IL Split | AAGCATGTGCAGTGCCTTC | Amplification of sulphonylurea resistance gene *ILV1* |
| LV1Split | CGCCCGGCCGACATCC | Amplification of sulphonylurea resistance gene *ILV1* |
| GRASPKOConF | TGTTGCCCTTGCCTTTCT | Confirmation of *MoGRH1* deletion |
| GRASPKOConR | CCAGCAATTCAACGACGATAAC | Confirmation of *MoGRH1* deletion |
| pGTN-Conf-Fw | CACGTTGTCGCCTAACAGAT | Confirmation of genomic integration of the pGTN vector carrying the *GFP-ATG8* gene |
| ATG8-SEQ-R | GAGACAGCTTGATGCGCTTGC | Confirmation of genomic integration of the pGTN vector carrying the *GFP-ATG8* gene |
| PWL2 Conf Fw | CTTATTATGGTCCCGGGTGAT | Confirmation of genomic integration of the pBV591 or modified pBV591 vector |
| qGFPRe | TGTTGTGGCGGATCTTGAA | Confirmation of genomic integration of the pBV591 or modified pBV591 vector |

**Supplementary Table S2: Primers used in the study**
